# Community interactions drive the evolution of antibiotic tolerance in bacteria

**DOI:** 10.1101/2022.06.02.494585

**Authors:** Sivan Pearl Mizrahi, Akshit Goyal, Jeff Gore

**Affiliations:** Physics of Living Systems, Department of Physics, Massachusetts Institute of Technology, Cambridge 02139, USA

**Author notes:** Equal contribution (Sivan Pearl Mizrahi, Akshit Goyal).

**Keywords:** antibiotics, tolerance, evolution, community interactions, cross-protection

## Abstract

The emergence of antibiotic tolerance (prolonged survival against exposure) in natural bacterial populations is a major concern. Since it has been studied primarily in isogenic populations, we do not yet understand how ecological interactions in a diverse community impact the evolution of tolerance. To address this, we studied the evolutionary dynamics of a synthetic bacterial community composed of two interacting strains. In this community, an antibiotic-resistant strain protected the other, susceptible strain by degrading the antibiotic ampicillin in the medium. Surprisingly, we found that in the presence of antibiotics, the susceptible strain evolved tolerance. Tolerance was typified by an increase in survival as well as an accompanying decrease in growth rate, highlighting a trade-off between the two. A simple mathematical model explained that the observed decrease in death rate, even when coupled with a decreased growth rate, is beneficial in a community with weak protective interactions. In the presence of strong interactions, the model predicted that the trade-off would instead be detrimental and tolerance would not emerge, which we experimentally verified. By whole genome sequencing the evolved tolerant isolates, we identified three genetic hotspots which accumulated mutations in parallel lines, suggesting their association with tolerance. Our work highlights that ecological interactions can promote antibiotic tolerance in bacterial communities, which has remained understudied.

**Significance:** Bacteria evolve to evade antibiotic pressure, leading to adverse infection outcomes. Understanding the evolutionary dynamics which lead to different antibiotic responses has thus far focused on single-strain bacterial populations, with limited attention to multi-strain communities which are more common in nature. Here, we experimentally evolved a simple two-strain community, comprising an antibiotic-resistant strain protecting a susceptible one, and found that susceptible populations evolve tolerance, helping them better survive long antibiotic exposure. Using the interplay between community interactions, antibiotic dynamics, and resource availability, we explain this finding with a simple mathematical model, and predict and experimentally verify that an increased resistant strain carrying capacity would render tolerance detrimental. Our results highlight that community interactions can alter bacterial evolutionary responses to antibiotics.

## Introduction

The extensive use of antibiotics over the decades has brought with it a growing concern of the ability of bacteria to evolve and evade them. Bacteria can evolve a variety of strategies to survive antibiotics, by far the most familiar being resistance, the ability to grow in higher antibiotic concentrations. Another common, but less well-studied, bacterial strategy against antibiotics is tolerance — the ability to survive antibiotic exposure for longer duration (1). A plethora of studies have revealed the many mechanisms by which bacteria can evolve both resistance and tolerance, in clinical as well as laboratory conditions (2–5). However, laboratory studies, which provide tractability and control, have largely been limited to studying single strains of bacteria, isolated from the community context in which they naturally occur. In contrast, clinical studies provide natural context, but limit experimental control and tractability. To understand how community context shapes bacterial response to antibiotics, we need experimentally tractable engineered communities which can evolve, whose composition we can manipulate, and whose evolved members we can characterize.

Interactions between bacteria can profoundly shape the dynamics and structure of ecological communities, including but not limited to their growth in the face of antibiotic exposure. Indeed, antibiotic-resistant strains can protect susceptible strains against antibiotics, allowing the two to coexist even in high antibiotic concentrations, far exceeding the minimum inhibitory concentration (MIC) of the susceptible cells (6–8). Interactions in a community have also been shown to change the sensitivity of a focal strain (9) or impact its tolerance (10). Further, competition and cooperation for nutrients could also affect community dynamics, and in turn the costs and benefits of resistance and tolerance (11–13). Community context has been shown to impact the evolution of resistance in general in a variety of environmental contexts, such as sewage (14) and animal feces (15). Finally, *β*-lactamase producing strains were shown to impact the course of poly-microbial infections (16). Thus, both the type and strength of inter-species interactions can shape the fitness of antibiotic-tolerant as well as resistant bacteria, and modify their evolutionary trajectories.

Here, we studied the evolutionary dynamics of a simple synthetic two-strain bacterial community exposed to the antibiotic ampicillin, where a resistant strain degraded ampicillin in the medium, resulting in the protection of a susceptible strain. Surprisingly, we observed that the susceptible strain repeatedly evolved tolerance in multiple parallel lines of this community, typified by a reduced death rate and a concomitant decrease in growth rate. By mathematically modeling this community, we could explain why tolerance was beneficial in this community context, and predicted that tolerance would become detrimental if we increased the protection provided by the resistant strain. Finally, we experimentally manipulated the strength of protection in our engineered community, and found that tolerance did not emerge in these conditions, as predicted by our model. Taken together, our results highlight the importance of studying antibiotic response through the lens of community interactions.

## Results

### Antibiotic exposure in communities leads to the evolution of tolerance

To understand how antibiotic responses evolve in a community context, we assembled and experimentally evolved a synthetic community of two *E.coli* strains, one susceptible to ampicillin and the other resistant to it (Fig. 1a, Methods). The resistant strain was an auxotroph for lysine and could not grow well without it being supplemented to the growth medium (Fig. S1). The auxotrophy of the resistant strain allowed us to control its population size by supplementing different amounts of lysine, and thus (as we later show) tune the interaction between the resistant and susceptible strain. Both strains could be distinguished by color and selective plating (see Methods). When the susceptible strain was exposed in monocultures to ≥ 64*μ*g/mL ampicillin (over 50-fold the MIC, 1 *μ*g/mL), the population rapidly died and went extinct within 3 daily dilution cycles into fresh media supplemented with ampicillin. However, the susceptible population could survive similar ampicillin exposure and stably coexist in a community alongside resistant cells in the presence of similar daily growth and dilution cycles (Fig. 1c).

**Figure 1.**
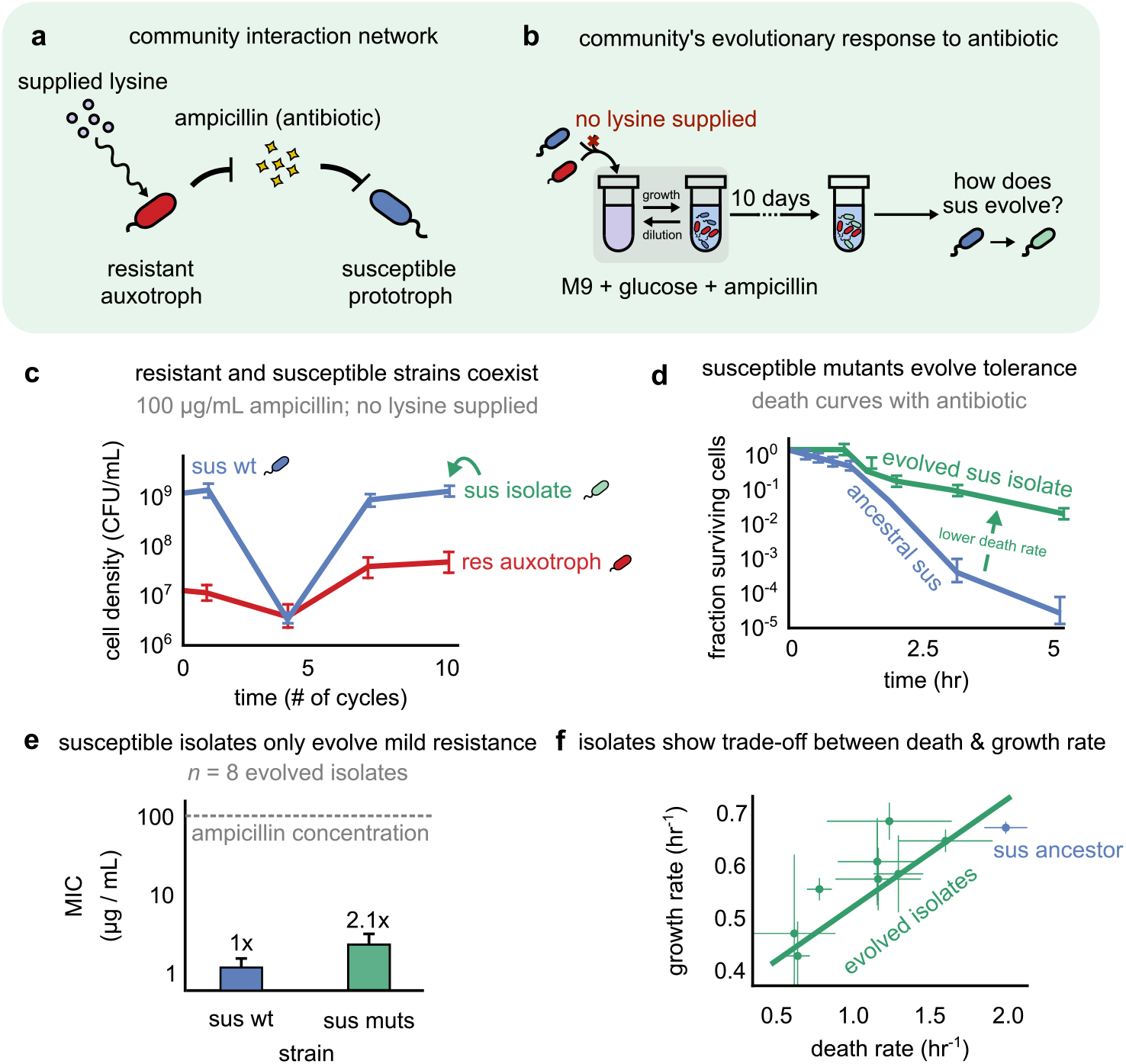
In an interacting community with protective interactions, antibiotic exposure leads to the evolution of tolerance. (a) Community interaction network, where a resistant strain (red), auxotrophic for lysine (circles), degrades ampicillin (diamonds), in turn protecting a susceptible prototroph (blue). By supplementing lysine, we can modulate the carrying capacity of resistant auxotrophs (curly arrow). (b) Schematic showing our experimental protocol, where we mixed susceptible (sus) and resistant (res) strains with no lysine, and subjected them to 10 cycles of growth and dilution. At the end of 10 cycles, we isolated susceptible strains and asked what antibiotic response they had evolved. (c) Abundance over different cycles of both strains in the community during an evolutionary experiment. Values show the mean of multiple replicates plating for one representative line. (d) Kill curves of the isolated ancestral strain (sus wt, blue) and one representative evolved isolate (green), in 100 *μ*g/mL ampicillin. Values represent the mean across technical replicates. (e) Bar graph showing the average minimum inhibitory concentrations (MIC) of the susceptible wild-type (ancestor) and eight evolved isolates, Mann-Whitney U test,two-sided, *p* < 0.005. (f) Scatter plot showing the death rate and growth rate for the ancestor (blue, *n* = 8) and evolved isolates (green, *n* = 8) from the evolutionary experiments as described in 1b. Line denotes the linear fit with *R*^2^ = 0.84 and *p* < 0.001. Values show mean of biological repeats. In all cases, error bars show s.e.m.

In this community, resistant cells protected susceptible cells by secreting the enzyme β-lactamase and degrading ampicillin, such that ampicillin levels dropped below the minimum inhibitory concentration (MIC) of the susceptible cells (Fig. S4), as previously reported for a similar community (6). We found only limited evidence of the susceptible (lysine prototroph) strain benefiting the resistant (lysine auxotroph) strain (Fig. S3). Since the susceptible strain could survive ampicillin exposure in such a community setting, we hypothesized that it might evolve to better respond to the repeated ampicillin exposures. In particular, we were interested in understanding how the susceptible strain would evolve in response to antibiotic exposure (Fig. 1b); whether it would increase its MIC (gain resistance), as naively expected, or evolve an alternate strategy, say elongating the duration it can withstand death due to antibiotic exposure (increased tolerance).

To characterize the evolutionary response of the susceptible strain upon ampicillin exposure within our communities, we performed experimental evolution under high ampicillin concentrations, far exceeding the MIC of the susceptible strain, but much below the MIC of the resistant strain (see Methods). Each day the co-cultures were diluted 1:50 into fresh media with high ampicillin (≥ 64× the MIC). We continued to serially propagate the communities for 10 growth-dilution cycles and measured the abundance of the two strains periodically. We did not study cases where the susceptible population went extinct, presumably due to stochastic evolutionary dynamics (Fig. S2 for statistics). At the end of 10 cycles (approximately 100 generations), we isolated representatives of the evolved susceptible strains.

Surprisingly, while the evolved isolates showed only a moderate increase in their MIC (from 1 *μ*g/mL to ~2*μ*g/mL; Fig. 1e), they had evolved much lower death rates (average decrease of 48% compared to wild type isolates; Fig. 1d,f). To test if the growth of the evolved isolates had been affected during evolution as well, we measured their growth rates. Interestingly, we found that the isolates also had reduced growth rates, and that the decrease in growth rate was linearly proportional to the decrease in death rate (Fig. 1f). The observed increases in MIC, indicative of resistance, were far from the exposed antibiotic concentrations (100 *μ*g/mL). Moreover, a two-parameter regression showed that the increased survival of evolved isolates, quantified by their surviving frequency after 5 hours of antibiotic exposure, was much better explained by the decrease in growth rates than increases in the MIC (regression coefficients 3.2±0.2, *p* 10 < ^3^ for growth rate versus −0.06±0.04, *p* = 0.02 for MIC). This suggested that the increase in MIC alone was not sufficient to explain the increased survival of the evolved isolates; instead, the increased survival was likely due to tolerance. Thus, we concluded that to respond to antibiotics in our synthetic communities, susceptible strains evolved tolerance by slow growth i.e., a decrease in death rate, coupled with a decrease in growth rate (17).

### A mathematical model explains that tolerance is beneficial under weak protective interactions

Having learnt that tolerance repeatedly evolved in our susceptible populations, we sought to understand what makes this strategy beneficial. In principle, a decreased death rate should always be advantageous, since it helps more susceptible cells survive when antibiotic levels are above the MIC. However, the tolerance observed in our populations has both a benefit (decreased death rate) as well as a cost (an accompanied decrease in growth rate). Thus, we would only expect tolerance to be beneficial in conditions when the benefit outweighs the cost.

To understand when a decreased death rate is beneficial despite an accompanying decreased growth rate, we calculated the relative fitness of tolerant isolates in different conditions, using a simple mathematical model of our synthetic microbial community. Briefly, our model simulated the growth of a community with three distinct strains: resistant auxotroph which grew at a fixed growth rate *γ_R_* regardless of the antibiotic concentration (Fig. 2a, red), susceptible ancestor which grew at a rate *γ_S_* at antibiotic concentrations below the MIC, and died at a rate *δ_S_* when antibiotic concentrations are above the MIC (Fig. 2a, blue), and finally, a tolerant isolate of the susceptible strain, with a reduced growth rate *γ_M_* and death rate *δ_M_* (Fig. 2a, green). We assumed that the decrease in death and growth rates were linearly related, consistent with our experimental observations (Fig. 1f,Fig. 2a, bottom; Methods) as well as previous findings (18,19). The entire bacterial community was subjected to daily dilutions, mimicking our experimental protocol (details in Methods). In the first growth-dilution cycle we seeded each community with a certain number of each strain (See Methods). Each community was subjected to an externally controlled antibiotic concentration at the beginning of each day. During each cycle, after an initial one hour lag, resistant auxotrophs grew at a rate *γ_R_*. The rate of antibiotic degradation was proportional to the number of resistant auxotroph cells present, regardless of their instantaneous growth rate. After a similar one hour lag, susceptible and tolerant populations both declined at their respective death rates until the antibiotic concentration dropped below the MIC, after which they grew at their respective rates. To model the lysine-dependence of resistant auxotrophs, we controlled their carrying capacity *R_sat_*; increasing lysine concentrations corresponded to an increasing *R_sat_* (Fig. 2b and 2d; Methods). For simplicity, the community had a fixed carrying capacity, *N_sat_*; when the community size hit *N_sat_*, growth ceased until the next cycle (Fig. 2a, middle).

**Figure 2.**
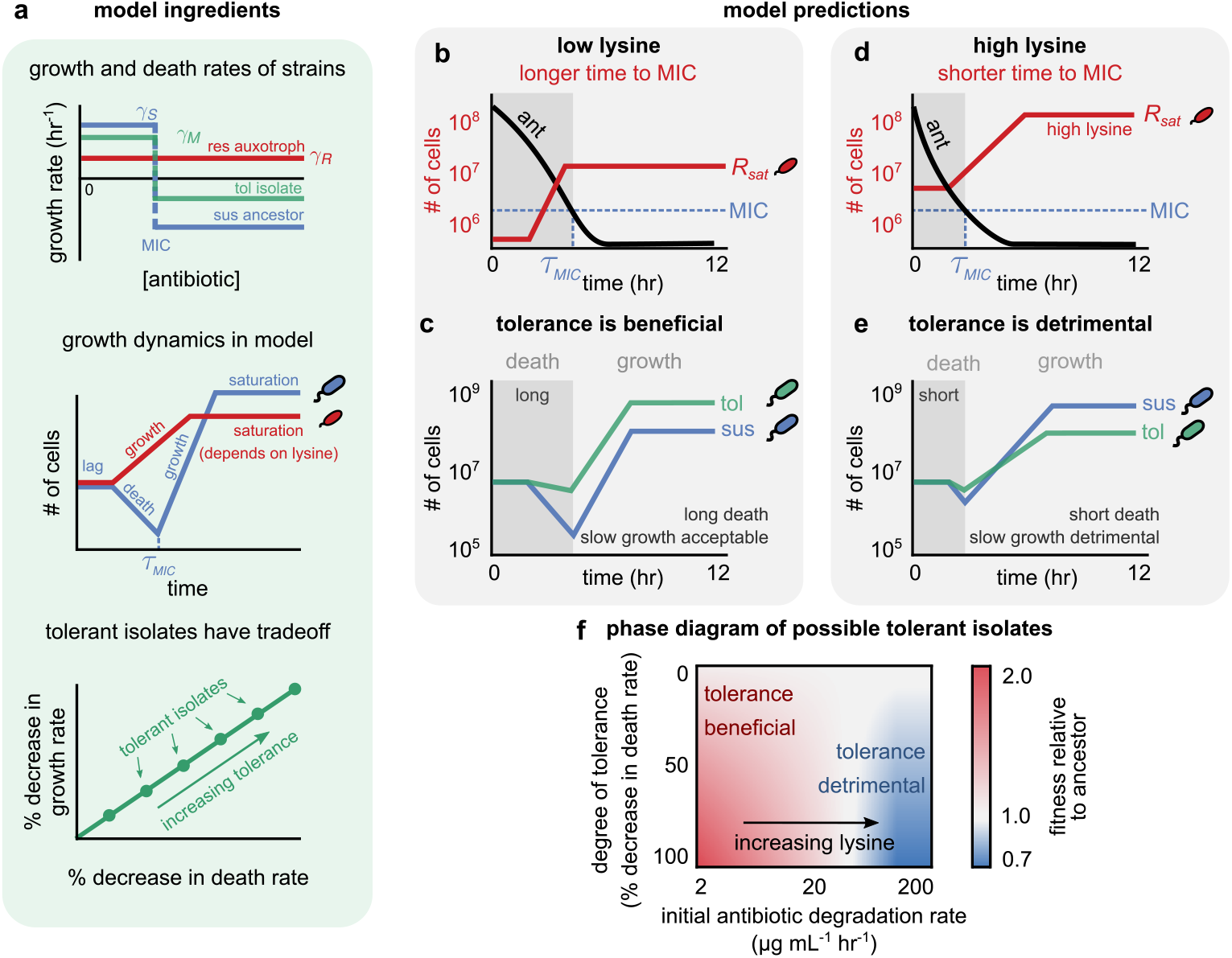
A mathematical model explains that tolerance is only beneficial when the community has weak protective interactions. (a) Ingredients of our mathematical model. Top: plot showing the growth rate as a function of antibiotic concentration for three strains: the resistant auxotroph (red), susceptible ancestor (blue), and a tolerant mutant (green). Middle: example of dynamics of resistant (red) and susceptible (blue) strains. The resistant strains grow till they hit their carrying capacity, the parameter *R_sat_* in our model, which is analogous to the lysine concentration in our experiment. Bottom: scatter plot showing that all tolerant strains in the model obey a linear relationship between their growth and death rates. (b–e) Dynamics of resistant strains (red), the antibiotic (black), susceptible ancestor (blue) and tolerant mutant (green) under (b–c) low lysine and (d–e) high lysine concentrations (in our model, low *R_sat_* and high *R_sat_*). The dynamics are a cartoon representation (see Fig. S12 for simulated dynamics); for how we measure the relative fitness, see Methods. The horizontal dashed line shows the MIC of the susceptible ancestor, and the grey shaded region shows the region where the antibiotic concentration is above the susceptible strain’s MIC. (f) Heatmap showing the relative fitness of possible tolerant strains in the model with varying degrees of tolerance and antibiotic degradation rates (controlled by changing the resistant population size).

Simulations of our model suggested that tolerance is beneficial when the population size of resistant auxotrophs is small, resulting in slow (~ 5 hrs) antibiotic degradation and weak protection of susceptible cells, i.e., a long time for the antibiotic to drop below susceptible cells’ MIC (Fig. 2b, black). In our model, this occurs when the resistant auxotroph population has a low carrying capacity, which we experimentally achieved by not supplementing lysine. In these conditions, susceptible strains face a long death phase (Fig. 2c, grey region), during which their population declines significantly (Fig. 2c, blue). Thus, tolerant isolates, which have a lower death rate, are fitter, even at the cost of a lower growth rate (Fig. 2c, green). Since tolerant populations decline significantly less than susceptible populations, once they start growing, they can divide more than their ancestors in a single cycle, making them fitter (See Supplementary text). The model also explained that the benefit of tolerance depends on the slope of the growth-death trade-off line (Fig. 1f and Fig. 2a, bottom). Namely, if the slope of the trade-off is too steep, such that a small reduction in death rate results in a large reduction in growth rate, then tolerant isolates become relatively less fit. Specifically, there is a threshold slope of this line above which tolerance ceases to be beneficial (Supplementary Fig. S5). Moreover, the degree of tolerance also depends on the antibiotic concentration — in lower concentrations, where susceptible death phase is shorter, a large decrease in death rate is not beneficial (Fig. S6). Finally, increase in MIC only weakly contributed to the relative fitness of tolerant strains in the model (Fig. S7). Specifically, increasing the MIC of tolerant strains in the model to 2 *μ*g/mL (the MIC measured in our experimentally evolved isolates) increased the relative fitness of tolerant strains by less than 10%. Thus, the model suggested that tolerance is the dominant contributor to the fitness of our evolved isolates, not resistance.

Taking the results from the model together, we concluded that when the resistant cells offer weak protection from the antibiotic in the form of slow degradation, to the susceptible cells in the community, we should expect tolerance to emerge, as seen in our evolved isolates.

### Tolerance does not evolve in communities with faster antibiotic degradation

Having understood that evolving tolerance could make a susceptible population more fit in a community with slow antibiotic degradation, we next asked in which conditions tolerance would not be favored. Interestingly, our model predicted that strengthening protection from the resistant auxotroph — by increasing its carrying capacity and thereby speeding up antibiotic degradation — will abolish the benefits of tolerance by slow growth. This is because tolerant isolates are relatively less fit than their susceptible ancestors in these conditions. With a higher carrying capacity, a larger resistant auxotroph population degrades the antibiotic much faster. Due to this faster degradation, susceptible strains spend a shorter period in the death phase compared with conditions with slow antibiotic degradation (Fig. 2d); instead they spend the majority of each cycle in the growth and saturation phases (Fig. 2e, grey, Fig. S4). It is ultimately because of a much shorter death phase that tolerant isolates are less advantageous than their susceptible ancestor (Fig. 2e, white). Using our model, we charted the phase space of tolerance evolution, plotting the relative fitness of tolerant isolates versus their susceptible ancestors under a wide range of parameters (Fig. 2f). Using this phase diagram, we concluded that tolerance is likely to be observed only when the decrease in growth rate of emerging tolerant isolates is not very large, and importantly, when antibiotic degradation is slow. A corollary of this was the prediction that tolerance would not emerge in conditions with rapid antibiotic degradation, achieved when the resistant cells had a large population size and strongly protected susceptible cells.

To test this prediction of our model, we repeated our evolution experiment, supplementing the growth medium with increasing concentrations of lysine, increasing resistant strain capacities (Fig. 3a, Fig. S8). In these conditions, the susceptible and resistant strains still coexisted (Fig. 3b). In agreement with our prediction, above 0.0001% lysine we didn’t observe any evolution of tolerance in 8 out of 9 lines (Fig. S9 and Fig. 3c-d). Susceptible cells isolated after 10 days of evolution showed neither a decrease in death rate (Fig. 3d), nor an increase in MIC (Fig. 3e). Indeed, our model predicted that when the antibiotic degradation rate was very large, such that there was no death phase for susceptible cells, there would be no advantage of having an increased MIC, even if this increase had no associated cost (in the model, this happens when *τ_MIC_* < *τ_lag_*, see Supplementary text). These results suggest that the evolution of tolerance can be suppressed in communities which strongly protect susceptible strains from antibiotics.

**Figure 3.**
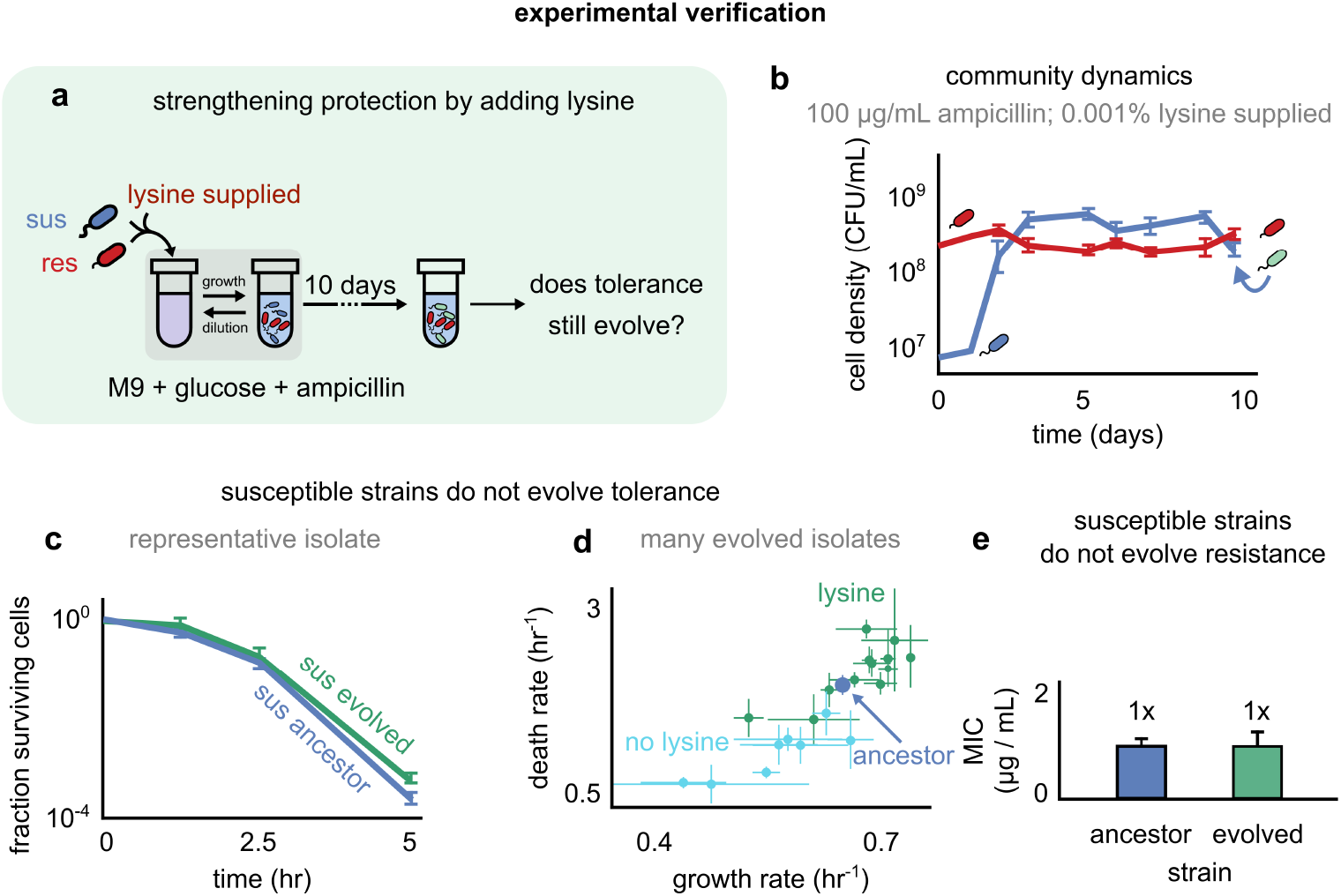
Strengthening community interactions (supplying lysine) during antibiotic exposure suppresses the evolution of tolerance. (a) Schematic showing our experimental protocol, where we mixed susceptible and resistant strains with lysine, and subjected them to 10 cycles of growth and dilution. At the end of 10 cycles, we isolated susceptible strains and asked which form of antibiotic response they had evolved. (b) Susceptible and resistant strains co-exist at high ampicillin exposure and when supplemented with lysine (0.001% lysine shown). Line plots showing the cell densities of the susceptible (blue) and resistant (red) strain at the end of each growth-dilution cycle, for a representative evolutionary experiment. (c) Kill curves for the susceptible ancestor (blue) as well as the evolved isolate (green) from the representative line in (b). Shown are the mean of technical replicates; error bars represent s.e.m. (d) Scatter plot showing the growth and death rates of different isolates evolved under lysine supplementation (green) and without lysine supplementation (cyan). The growth and death rate of the ancestral strain (anc) is marked in blue. Values and error bars represent the mean and s.e.m. (*n* = 8 for isolates evolved under no lysine and n = 12 isolates evolved under ≥ 0.0001% lysine). (e) Susceptible strains evolved in lysine concentrations ≥ 0.0001% (green, *n* = 12) had similar MIC compared with the ancestor (blue, *n* = 8);, Mann-Whitney U test, two-sided, p > 0.1.

Another way to shorten the duration over which susceptible cells die (lower *τ_MIC_*) is to reduce the initial antibiotic concentration. We repeated our evolutionary experiment using a lower ampicillin concentration of 15*μ*g/mL. Interestingly, susceptible strains evolved under low lysine and low ampicillin concentrations showed a lower reduction in death rates compared to those evolved under high ampicillin concentrations (average death rates of 1.2 ± 0.1 hr^−1^ versus 1.8 ± 0.1 hr^−1^; Mann Whitney one-sided test *p* < 0.05; Fig. S10, left panel). This difference was not significant for isolates evolved under high lysine (average death rates of 2.2 ± 0.1 hr^−1^ versus 2.0 ± 0.1 hr^−1^; Mann Whitney one-sided test p > 0.1; Fig. S10, right panel). These results suggest that the exposed antibiotic concentration can also control whether tolerance evolves and to what degree it evolves.

### Two classes of mutations are associated with tolerance by slow growth

To understand the genetic basis of the “tolerance by slow growth” observed in our study, we sequenced the whole genomes of selected evolved isolates. We sequenced all isolates with increased survival and a few with no increased survival to serve as controls. We identified a total of 12 isolates carrying mutations (Table S2). Our analysis revealed two genetic “hotspots” that had accumulated mutations in several parallel tolerant evolved lines: the *envZ* gene and the *gln* operon (Table S2). Of the 9 tolerant isolates with decreased death rates, 8 had identified mutations in these “hotspots”. Isolates with mutations in envZ genes had similar decreases in death and growth rates (Fig. 4, red). In contrast, out of the 10 isolates with no significant change in growth rates (Table S2), only 3 had any detectable mutations outside of the 2 “hotspots” (Fig. 4).

**Figure 4.**
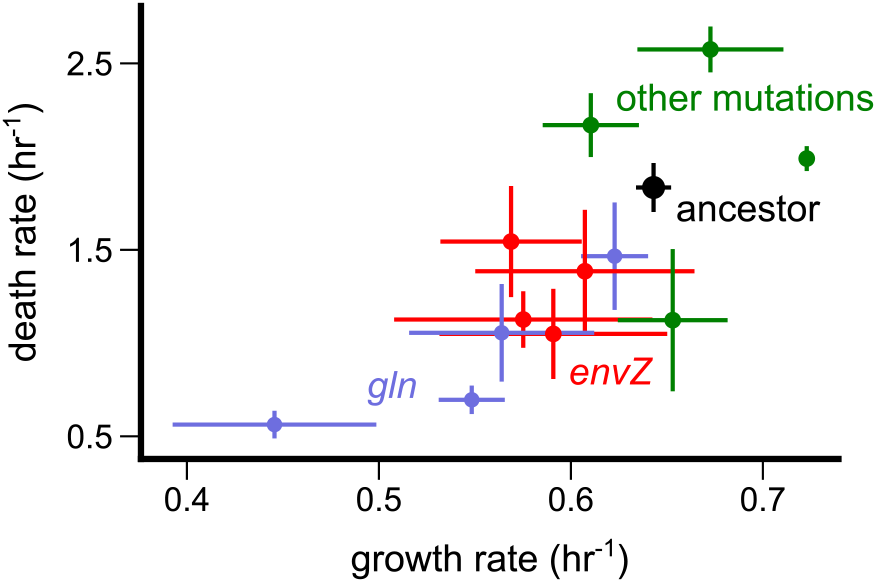
Classes of mutations associated with tolerance by slow growth. Death and growth rates of sequenced evolved isolates in which we identified mutations are plotted as well as the ancestral strain (black). Each point represents a distinct isolate, whose color represents which genes or pathway were mutated in it (*envZ* in red, *gln* in blue, other mutations in green). Points represent mean values, while error bars represent the s.e.m.

Previous literature suggests that these two genetic hotspots might be linked to the observed slow growth (tolerance). The *envZ* gene, in which we detected mutations in four parallel lines, is a membrane-associated protein involved in osmoregulation. Its expression has previously been linked to β-lactam resistance either by down-regulating *ompC* and *ompF* (20), or by a different pathway (21). Genes in the *gln* operon — namely *glnA, glnB* and *glnD* — are known for sensing, regulating, and metabolizing nitrogen. Nitrogen starvation, in part controlled by *gln* genes, can impact cell growth (22, 23). Interestingly, we also detected mutations in ribosomal RNA genes in parallel lines — in particular mutations in the *rrl* operon. Ribosome synthesis is a key growth-determining process in bacteria and mutations lowering its rate could simultaneously reduce both bacterial growth and death rates (24). However, tolerant isolates carrying rRNA mutations also had other mutations (Table S2), making it difficult to pinpoint any causal link between rRNA mutations and tolerance (25,26).

In conclusion, we discovered two genetic pathways associated with the emergence of tolerance. In our synthetic community, where resistant cells weakly protected susceptible cells, these mutations were associated with an increased fitness of susceptible populations.

## Discussion

In this study, we have shown that when exposed to antibiotics in a community context in which resistant bacteria protect susceptible ones, the susceptible population can evolve tolerance. This tolerance is characterized by a decrease in both the growth and death rates of the bacterial population, which improves their fitness in the community. Moreover, we showed that the evolution of tolerance can be suppressed if we modulate the interactions between resistant and susceptible bacteria, e.g., by increasing the carrying capacity of the resistant strain, which speeds up antibiotic degradation. We achieved this by experimentally evolving a synthetic community in which we could control the environmental conditions, antibiotic concentration, and the strength of interactions between resistant and susceptible bacteria. Finally, using a simple mathematical model of the community, we could also explain the costs and benefits of tolerance by slow growth in various conditions, and successfully predict under which conditions it would be expected to emerge in our experimental communities.

The strikingly linear relationship between the death rates and growth rates of our evolved isolates hints at a possible causal link between the two, perhaps even indicating an important biological trade-off. While our study did not focus on these possible links, future studies discerning the mechanism behind this pattern are likely to shed light on both the mechanisms and fundamental constraints driving such evolution in response to antibiotics in bacteria.

The evolution of mechanisms other than resistance has been observed in other studies with single species populations exposed to antibiotics above their MIC. One study exposed *E.coli* populations to different antibiotics during the stationary phase and reported the emergence of persistence (a larger sub population of tolerant cells) rather than resistance (27). Another experimental evolutionary study exposed *E.coli* populations to different antibiotics during the exponential phase, and found single point mutations leading to tolerance (to the specific drug class given) (28). Interestingly, the line evolved under repetitive ampicillin exposure, had a significant decrease in its growth rate. Under the conditions of this study persistence also evolved in all treatments while resistance did not evolve. In yet other studies, tolerance by lag evolved (4,29). One crucial difference between these studies and ours is that in our study, the antibiotic concentration is changed naturally by the community itself, instead of having it changed by the experimentalist. In this way, the community changes its own antibiotic landscape over time, and these interactions can make tolerance more beneficial than resistance.

While we focused on the evolution of the susceptible strain in our study, it is possible that the resistant auxotroph evolved in our experimental conditions as well. Indeed it was shown that strains in an obligate cross-feeding system, when exposed to antibiotics, evolved autonomous metabolic activity and weakened the mutualistic interactions (12). Moreover, the interactions between the two strains in our experiment might change during their evolution. Studying such changes over longer evolutionary timescales is, in our opinion, likely to be a fruitful avenue for future work.

Our model included several simplifying assumptions. First, as a proxy for the lysine concentration, we tuned the carrying capacity of the resistant strain, *R_sat_*. We did not explicitly model lysine, its dynamics, and dependence on the abundance of the susceptible strain (say in the form of it producing and secreting lysine). Second, we assumed that lysine concentration only affected the resistant strain’s carrying capacity and not its growth rate, supported by our mono-culture measurements (Fig. S1). However, in some co-cultures evolved under high ampicillin and high lysine, resistant cells survived while the susceptible strain went extinct (Fig. S2), which does not agree with our model. This could be recapitulated in a variant of our model where the growth rate of the resistant auxotroph increased with the lysine concentration. Third, we assumed that the resistant strain had an arbitrarily large MIC, not dying at any ampicillin concentration. Measurements of the resistant strain “single-cell MIC” (30) indeed show it is higher than 800*μ/mL* ampicillin. Moreover, we assumed that bacterial growth was not impacted by antibiotic concentrations below the MIC.

During the evolutionary experiment antibiotic concentrations drop continuously, exposing the susceptible strain to sub-MIC concentrations. Sub-MIC concentrations can lead to the evolution and spread of resistance (31,32). Moreover, there is an interplay between resistance and tolerance; indeed, tolerance has already been identified as a stepping stone for resistance (29,33,34). We observed a mild increase in the MIC in several of the isolates, albeit with a minor contribution to fitness when compared with tolerance. Nonetheless, in our model, any increase in MIC is beneficial, since it reduces the duration of the death phase (in the model, *τ_MIC_*) without any associated cost.

To summarize, we have shown that in a synthetic community in which an antibiotic resistant strain protects a susceptible strain, the evolution of tolerance is restricted by the strength of the protective interaction. This highlights the importance of expanding our knowledge of the intricate interactions between ecology and evolution, especially in the context of antibiotic treatments and its alarming increased failure.

## Materials and Methods

### Strains and Media

All strains are derived from Escherichia coli BW25113. The resistant auxotroph strain is based on JW2806-1 taken from the Keio collection (35) and is deleted for *lysA* and carries a kanamycin resistant gene. It contains the pFPV-mCherry plasmid (36) (also see Addgene plasmid 20956), expressing a *β*-lactamase enzyme and an mCherry fluorescent protein. The susceptible strain was constructed by P1 transduction of the *yfp-Cam* cassette from M2(37) to BW25113.

M9 media (Sigma cat# M6030) supplemented with 0.5*μ* g/mL B1 vitamin and 0.2% glucose was used for growth and in the evolution experiments. M9 was either without or supplemented with different concentrations of L-Lysine (Sigma cat# L5501) as indicated in the text. LB-agar plates were used for counting CFUs. Though the colonies of both strains could be differentiated by their colors, plates were frequently supplemented with either Chloramphenicol (25*μ*g/mL, Sigma cat# C0378) or Kanamycin (50*μ*g/mL,Sigma cat# K0254) to select for the susceptible and resistant strains respectively.

### Evolutionary protocol of daily exposure to antibiotics

Evolution under antibiotic exposure started with a 1:1 ratio of overnight susceptible and resistant strains diluted 1:50 into fresh media supplemented with ampicillin (Millipore cat# 171254) and lysine (as indicated in the text), volume indicated in Table S3. in each cycle, samples were incubated at 37°C with shaking at 300 r.p.m. for 24 hours and daily diluted 1:50 into fresh media. Samples were periodically plated on LB-agar plates (with and without antibiotic selection) at the end of cycles to monitor the capacities of both strains.

### Estimation of population densities (CFU/mL)

For CFU counting, two methods were used, depending on the number of samples monitored. For few samples the classical CFU of serially dilution and plating with glass beads on LB-agar plates was used. When many samples were monitored a 96-well plate was used. 10 *μ*L droplets of PBS-diluted samples were plated on 150-mm-diameter LB-agar plates with or without antibiotic selection. To prepare the 10 μL droplets, we serially diluted the experimental cultures via 10-fold dilutions (maximum dilution factor was 10^7^) with a Viaflo 96-well pipettor. Droplets were then transferred to the agar plates and allowed to dry, followed by incubation at 37 °C for a day until colonies were visible. The different dilutions allowed us to find a dilution at which colonies could be optimally counted and several plating replicates per isolates were performed to increase accuracy at measuring population densities. Resistant and susceptible strains were distinguished either by plating on LB-agar plates and counting red (resistant) and white (susceptible) colonies. This did not always give the best separation since when one population was much denser it obscured the other one. In these cases (and always for lower dilutions) we plated the samples on LB-agar plates with antibiotic selection; kanamycin and choloramphenicol were used to assess the density of the resistant and susceptible populations respectively.

### Antibiotic survival assays

To measure the survival under antibiotic treatments, overnight cultures, each grown from a single colony, were diluted 1:50 in fresh medium supplemented with (100 *μ*g/ mL) ampicillin. At indicated time points, aliquots of the cultures were sampled and CFUs/mL were measured.

### MIC assays

Each column in a 96-well plate was filled with fresh M9 +0.2% glucose supplemented with increasing amounts of ampicillin and inoculated with approximately 10^5^ bacteria per well. The plate was incubated overnight at 37°C with shaking at 300 rpm. The Minimum Inhibitory Concentration (MIC) was recorded as the highest concentration which supported growth (measured by OD_630_).

### Growth rate measurements

The growth rates of the ancestral and evolved strains were measured by monitoring *OD*_630_ in 96-well plates with the multi-well reader Infinite (Tecan, Switzerland) at 37°C with shaking. The growth rates were extracted by fitting the exponential part of the growth in 3 replicates for each strain.

### Description of the mathematical model

We formulated a mathematical model of our community, comprising three kinds of strains: a resistant auxotroph, a susceptible ancestor and an evolved tolerant strain. All strains differed from each other in their growth rates and death rates. The resistant auxotroph had a fixed growth rate *γ_R_* regardless of antibiotic concentration. The susceptible ancestor had a growth rate *γ_S_* below its MIC (fixed at 1 *μ*g/mL), and a death rate *δ_S_* above the MIC. To model the cost of resistance, we assumed that *γ_R_* < *γ_S_* by about 15%, similar to what we measured experimentally for our strains. The tolerant strain, which we assumed evolved from the susceptible strain, had reduced growth and death rates *γ_M_, δ_M_*, respectively; it had the same MIC of 1*μ*g/mL as the susceptible ancestor. To mimic our experimental observation, we assumed that the growth rate *γ_M_* and death rate *δ_M_* of any evolved tolerant strain were linearly related by a trade-off through the following equation:

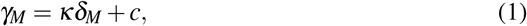

where *κ* represents the slope of the growth-death trade-off, and *c* represents the intercept. We fit *κ* = 0.21 and *c* = 0.75 hr^−1^ to match the experimentally observed slope and intercept, respectively (Fig. 1f). Using our model, we wished to study and compare the relative fitness of evolved tolerant strains with different values of *γ_M_* and *δ_M_*, all of which obeyed this trade-off relationship.

Using our model, we simulated the dynamics of all three strains growing together in a com-munity, undergoing several growth-dilution cycles. At the beginning of the first growth cycle, we added the resistant auxotroph and susceptible ancestor strains at equal population sizes at their steady state values, divided by the dilution factor *D*, and 1 cell of the evolved tolerant strain (to mimic an emerging mutant), as well as a particular concentration of antibiotic (typically 100 *μ* g/mL). Resistant cells degraded this antibiotic at a rate proportional to their current population size *R*, according to the following Michaelis-Menten dynamics for the antibiotic *A*:

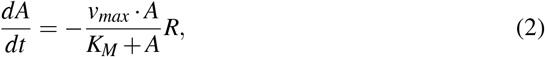

where *v_max_* was the maximal degradation speed and *K_M_* was the half-saturation constant of antibiotic degradation, respectively (parameter values matched to those observed). At the beginning of the growth cycle, all three strains experienced a lag period lasting 1 hour, approximately equal to that of the strains in our experiment. After this lag, the resistant cells *R* grew at a rate *γ_R_* according to the following equation:

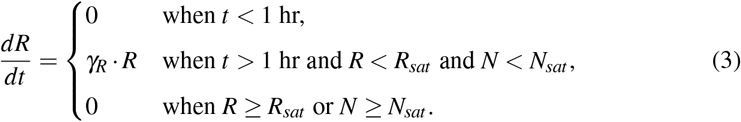

Here, *R_sat_* is the carrying capacity of the resistant cell population, which is a tunable parameter in our model, and *N_sat_* is the carrying capacity of the entire community, which we fixed to roughly 10^9^ cells. *N* = (*R* + *S* + *M*) represents the combined population of the entire community. We do not explicitly model lysine, and instead implicitly increase *R_sat_* to mimic an increase in supplemented lysine (as experimentally observed). The dynamics of the susceptible ancestor *S* and evolved tolerant strains *M* were as follows:

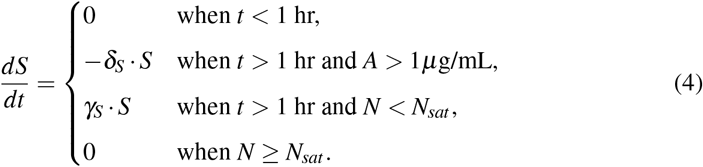

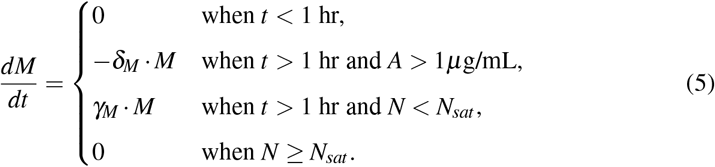

Each growth cycle lasted 24 hours, by which time the community population size reached *N* = *N_sat_*. After this 24 hour period, we diluted the community by a factor *D* = 50 (chosen to match the experiment), and began a new growth cycle with this new population in a medium replenished with the original antibiotic concentration. We repeated these growth-dilution cycles until the community reached steady state, that is, until all strains in the community reached the same population sizes at end of two consecutive cycles. We performed these simulations for each tolerant strain separately, where in each simulation we only added one tolerant strain at a small population size of 1 cell.

### Measuring the relative fitness of tolerant strains in the model

Using our model outlined above, we studied the benefits of evolving tolerance in different conditions — varying *γ_M_, δ_M_*, and *R_sat_* — by monitoring how much evolved tolerant strains grew compared with their susceptible ancestors in the same community. To quantify the benefits of tolerance, we measured the fitness *W* of a tolerant strain relative to its susceptible ancestor, defined as the ratio of the logarithms of their fold-growths over a growth cycle, as follows:

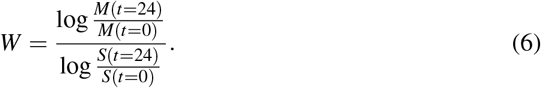

We were interested in the relative fitness of tolerant strains that emerged as mutants, and therefore measured this quantity in conditions where the susceptible ancestor and resistant auxotroph cells were in a steady state; in these conditions, the addition of 1 cell of a tolerant mutant would only negligibly affect their growth dynamics. In the Supplementary Text, we derive analytical expressions for the relative fitness as a function of the relevant dynamical quantities in our model.

### Sequencing and genomic analyses of isolates

We grew all strains from a single colony in 3mL LB at 37°C, overnight with shaking. We extracted genomic DNA using Zymo Quick-DNA plus miniprep kit (Cat No. D4068), and sent prepared samples of each strain to Quintara Biosciences for library preparation and Hi-Seqx2×150 sequencing. We trimmed all raw sequencing reads using Trimmomatic 0.36, using default settings. Using MiniMap2 v2.17, we then mapped all trimmed reads, sample by sample, to the *E. coli* reference genome. We used the reference genome for the strain BW25113, obtained from the NCBI RefSeq database. We edited this reference sequence to add an insertion sequence, whose details have been described in the section “Strains and Media”. In mapping reads to the reference genome, we ensured unambiguous read mapping by using the settings -`ax sr`. The resulting read alignments had an average coverage of 221. To identify mutations and genetic variation in the sequenced samples, we used the Bayesian genetic variant detector FreeBayes v1.3.2 using the following settings: --`ploidy 1 --haplotype-length 0 --min-alternate-count 1 --pooled-continuous`. For downstream analysis, we only used genetic variants (SNPs and indels) with a Phred quality score ≥ 30 and minimum local read depth 30. We analyzed the resulting set of variants that passed these filters. To map the variants with known gene annotations, we used the annotations for the strain BW25113 from the NCBI database, with the appropriate correction for our insertion sequence. For all samples, we filtered out all variants identified in the susceptible ancestor, compared with the BW25113 sequence, arguing that these mutations merely differentiated our laboratory strain from the reference sequence in the NCBI database.

## Acknowledgments

We thank NQ Balaban, O Gefen, I Levin-Reisman, O Fridman, S Moreno-Gaémez and N Shoresh for discussions. This work was supported by NIH Grant R01-GM102311 and the Schmidt Science Polymath Award. S.P.M. is supported by the HFSP LT000378/2018. A.G. is supported by the Gordon and Betty Moore Foundation as a Physics of Living Systems Fellow through grant number GBMF4513.

## Author contributions

J.G and S.P.M. designed the project. S.P.M. carried out the experiments. A.G and S.P.M. developed and analyzed the model. A.G performed the genomic analysis. J.G. supervised the study. All authors wrote the manuscript.

