## Supplementary Materials for "Community interactions drive the evolution of antibiotic tolerance in bacteria"

### Table S1: parameters used in the model

| Symbol | Meaning | Typical value (if applicable) |
|---|---|---|
| $\gamma_S$ | Growth rate of susceptible ancestor | 1.3 $hr^{-1}$ |
| $\delta_S$ | Death rate of susceptible ancestor | 2.7 $hr^{-1}$ |
| $\gamma_R$ | Growth rate of resistant auxotroph | 1.1 $hr^{-1}$ |
| $\gamma_M$ | Growth rate of tolerant evolved strain | Variable |
| $\delta_M$ | Death rate of tolerant evolved strain | Variable |
| $R_{sat}$ | Carrying capacity of resistant auxotroph | Variable |
| $\tau_{lag}$ | Duration of lag period in a growth cycle during which no strains grow | 1 hr |
| $v_{max}$ | Maximum degradation rate of antibiotic per unit antibiotic concentration | $10^{-5}\mu$g/cell/s |
| $K_M$ | Half-saturation constant for antibiotic degradation | 6.7 $\mu$g/mL |
| $N_{sat}$ | Carrying capacity of total community | $1.2 \times 10^9$ cells |
| $A(t=0)$ | Initial antibiotic concentration at the beginning of each growth cycle | 100 $\mu$g/mL |
| $D$ | Dilution factor | 50 |

### Calculating the relative fitness of tolerant strains in the model

Using our model described in the main text, we wished to calculate the relative fitness $W$ of any possible tolerant mutant as a function of various parameters, such as the carrying capacity of resistant auxotrophs $R_{sat}$, tolerant mutant (M) growth and death rates $\gamma_M$ and $\delta_M$, as well as the antibiotic concentration $A$. The relative fitness $W$, defined as the ratio of logarithms of the mutant and ancestral strains' fold-growths over a growth cycle, can be written as follows:

$$W = \frac{\log \frac{M(t=24)}{M(t=0)}}{\log \frac{S(t=24)}{S(t=0)}} = \frac{-\delta_M(\tau_{MIC,M} - \tau_{lag}) + \gamma_M(\tau_{sat} - \tau_{MIC,M})}{-\delta_S(\tau_{MIC,S} - \tau_{lag}) + \gamma_M(\tau_{sat} - \tau_{MIC,S})}. \tag{1}$$

Here, $\tau_{MIC,M}$ and $\tau_{MIC,S}$ are the times taken for the antibiotic to drop below the MICs of the mutant (M) and susceptible ancestor (S), respectively; $\tau_{sat}$ is the time it takes for the entire community to reach saturation density $N_{sat}$; and $\tau_{lag}$ is a fixed lag time for all strains. Importantly, the fitness $W$ is density-dependent, and its value for the same set of strains will change from the first growth cycle where the mutant is introduced at 1 cell, to a possibly new steady state growth cycle where the mutant population might be substantial. To predict whether an emerging mutant will fix or go extinct in the community, we are interested in its relative fitness when rare, and thus henceforth calculate $W$ at the growth cycle where the resistant auxotroph and susceptible ancestor are in the steady state, and any mutant of interest starts at a population size of 1 cell. In this case, we can safely assume that the mutant's growth only negligibly affects the growth and dynamics of the other two strains.

To calculate $W$ in these conditions, we must calculate $\tau_{MIC,S}$, $\tau_{MIC,M}$ and $\tau_{sat}$ for the community with the resistant auxotroph and susceptible ancestor at steady state. We will now calculate expressions for these quantities.

#### Derivation of antibiotic degradation time, $\tau_{MIC}$

In our model, we assume that the antibiotic is degraded by the resistant auxotrophs at a rate proportional to their population size $R$, through the following equation:

$$\frac{dA}{dt} = -\frac{v_{max} \cdot A}{K_M + A} R, \tag{2}$$

By integrating this equation, we can obtain the time taken by the antibiotic to drop to any desired concentration, such as the MIC of the susceptible ancestor, $A_{MIC}$. We get the following expression:

$$(A + K_M \log A)\Big|_{A_{(t=0)}}^{A_{MIC}} = -v_{max} \int_0^{\tau_{MIC}} R(t)dt. \tag{3}$$

Here, $t = 0$ represents the beginning of the steady state cycle. We can evaluate the integral on the right in two ways, depending on whether the resistant cells reach their carrying capacity $R_{sat}$ (time taken: $\tau_R$) before $\tau_{MIC}$, or after. We will evaluate these cases now.

#### Case I: When $\tau_R < \tau_{MIC}$

In this case, resistant cells reach their carrying capacity $R_{sat}$ at time $\tau_R$, after which their population size remains fixed at $R_{sat}$. Accordingly, the integral in equation (3) becomes the following:

$$(A + K_M \log A)\Big|_{A_{(t=0)}}^{A_{MIC}} = -v_{max}\left[R_{sat}(\tau_{MIC} - \tau_R) + \frac{R_{sat}}{D \cdot \gamma_R} e^{\gamma_R(\tau_R - \tau_{lag})} - \frac{R_{sat}}{D \cdot \gamma_R} + \frac{R_{sat} \cdot \tau_{lag}}{D}\right], \tag{4}$$

where we have used the fact that, at steady state, $R(t=0) = \frac{R_{sat}}{D}$, $D = 50$ being the dilution factor in our study. Rearranging these terms, and using the relation $e^{\gamma_R(\tau_R - \tau_{lag})} = D$ (which implies $\tau_R = \tau_{lag} + \log D/\gamma_R$), we get the following expression for $\tau_{MIC}$:

$$\boxed{\tau_{MIC} = \frac{1}{\gamma_R}\left[\log D - \frac{D-1}{D}\right] + \tau_{lag}\frac{D-1}{D} - \frac{A_F}{R_{sat}\cdot v_{max}},} \tag{5}$$

where $A_F = (A + K_M \log A)\Big|_{A_0}^{A_{MIC}}$. Typically, $D \gg 1$, so this equation can be simplified as follows:

$$\tau_{MIC} = \frac{1}{\gamma_R}\left[\log D - 1\right] + \tau_{lag} + \frac{A_F}{R_{sat}\cdot v_{max}}. \tag{6}$$

**Case II: When $\tau_R > \tau_{MIC}$**

This case further breaks down into two sub-cases: one where the entire community reaches carrying capacity $N_{sat}$ before resistant cells reach their own carrying capacity $R_{sat}$, and the other where the community saturates after the resistant cells reach capacity.

**Case IIa: When $\tau_R > \tau_{MIC}$ and $\tau_R < \tau_{sat}$**

We will now solve for the case where the resistant cells reach carrying capacity before the community does, we will have a very similar expression to the one in equation (4), except in this case, the resistant cells will not saturate before $\tau_{MIC}$, and therefore we will have one less term. Following the same arguments as before, the integral on the left hand side of equation (3) will be evaluated as follows:

$$\boxed{A_F = -v_{max}\left[\frac{R_{sat}}{D\cdot\gamma_R}e^{\gamma_R(\tau_{MIC}-\tau_{lag})} - \frac{R_{sat}}{D\cdot\gamma_R} + \frac{R_{sat}\cdot\tau_{lag}}{D}\right],} \tag{7}$$

which we can use to solve for $\tau_{MIC}$.

**Case IIb: When $\tau_R > \tau_{MIC}$ and $\tau_R \geq \tau_{sat}$**

In the case where the community reaches its carrying capacity before resistant cells do. At equilibrium, all subpopulations obey the following relation:

$$\frac{N_{sat}}{N(t=0)} = \frac{R(t=24)}{R(t=0)} = \frac{S(t=24)}{S(t=0)} = e^{\gamma_R(\tau_{sat}-\tau_{lag})} = D. \tag{8}$$

Here, $\tau_{sat} = \tau_{lag} + \frac{1}{\gamma_R}\log\Big(D\Big)$. When $\tau_{MIC} > \tau_{lag}$, i.e., the susceptible cells experience a death phase, we can write the following expression:

$$\log\left(\frac{S(t=24)}{S(t=0)}\right) = \log D = \gamma_S(\tau_{sat} - \tau_{MIC}) - \delta_S(\tau_{MIC} - \tau_{lag}). \tag{9}$$

Substituting the expression for $\tau_{sat}$ into equation (9), and solving for $\tau_{MIC}$, we get the following:

$$\boxed{\tau_{MIC} = \frac{1}{\gamma_S + \delta_S}\left[(\gamma_S + \delta_S)\tau_{lag} - \frac{\gamma_S - \gamma_R}{\gamma_R}\log D\right].} \tag{10}$$

Notice that in each of these cases, $\tau_{MIC}$ always depends weakly on the dilution factor $D$, through a $\log D$ dependence. However, how it depends on the initial antibiotic concentration $A(t=0)$ and resistant carrying capacity $R_{sat}$ can change depending on case. In the first case, which is where our experimental communities typically reside, $\tau_{MIC}$ depends linearly on $A(t=0)$ and inversely on $R_{sat}$, explaining how increasing $R_{sat}$ in our model would tend to decrease $\tau_{MIC}$. In the second case associated with equation (7), there is a weak dependence on both $A(t=0)$ and $R_{sat}$, (because $\tau_{MIC}$ now depends on their logarithms instead) and this again would be more likely at high resistant carrying capacities (i.e., high lysine concentrations). In the final case where $R_{sat}$ is never reached, highlighted in equation (10), there is no dependence on either quantity, suggesting that changes in the antibiotic concentration or resistant carrying capacity are not likely to affect degradation time at steady state. Note that for this case to be valid, the resistant cells would have to continue growing throughout the steady state cycle, which would only be likely when their capacity $R_{sat}$ is large (i.e., at high lysine concentrations in our experiment). In this case, $\tau_{MIC}$ is usually small enough for tolerance to not be beneficial (i.e., $W < 1$).

##### Derivation of community saturation time, $\tau_{sat}$

**Case I:** $\tau_{sat} \leq \tau_R$

We already derived an expression for $\tau_{sat}$ when the community saturates before the resistant cells do, using the following steady state expression for the growth of resistant cells:

$$\gamma_R(\tau_{sat} - \tau_{lag}) = \log D, \tag{11}$$

which must be obeyed at equilibrium when resistant cells don't saturate at $R_{sat}$.

**Case II:** $\tau_{sat} > \tau_R$

For cases when resistant cells do saturate at $\tau_R$, we use the following expression for the growth of susceptible cells:

$$\log\left(\frac{N_{sat}}{N(t=0)}\right) = \log D = \gamma_S(\tau_{sat} - \tau_{MIC}) - \delta_S(\tau_{MIC} - \tau_{lag}). \tag{12}$$

We can use equation (12) to solve for $\tau_{sat}$, by substituting the expressions for $\tau_{MIC}$ we derived in the previous section (equations (6) and (7)), according to which case is valid.

##### Estimating the parameters $v_{max}$ and $K_M$

We assumed a fixed value of the antibiotic affinity $K_M$, obtained from [3] as $K_M = 6.7\mu$g/mL. To estimate $v_{max}$, we used the following expression relating the equilibrium composition of the community, which can be derived using the expressions in the previous sections:

$$f_{eq} = \frac{R_{sat}}{N_{sat}} = \frac{A(t=0)\gamma_R\gamma_S}{v_{max}\cdot(\gamma_S - \gamma_R)N(t=0)}. \tag{13}$$

By measuring the fraction of resistant cells relative to the susceptible cells at equilibrium in the community, we could plug in that value into equation (13), including all other parameters which were known or measured, and solve for $v_{max}$. We obtained a value of $v_{max} = 10^{-5}\mu$g /cell /s.

### Ribosomal RNA mutations identified in this study

Our genetic analyses revealed rRNA operon mutations that might be associated with tolerance by slow growth. While it is plausible to imagine a causal link between ribosomal mutations and growth rate, previous work has shown that deletions of rRNA operons do not typically affect growth rate in *E. coli* [2, 1]. Interestingly, all but one of our evolved isolates with rRNA mutations also had mutations in other loci, making it hard to pinpoint a causal relationship between the observed rRNA mutations and a reduced growth rate. Consistent with previous work, the one evolved isolate from our study whose only mutations were in an rRNA operon did not have a significantly different growth rate.

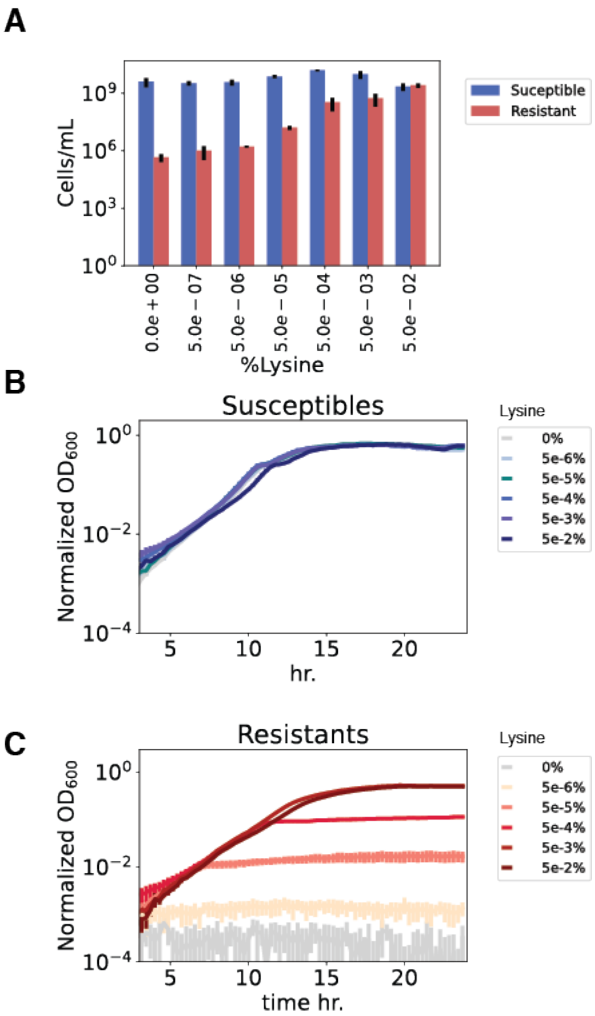


**Figure S1. Carrying capacity and growth curves of susceptible and resistant strains** (a) Cell densities of both strains after 24 hr of growth in media supplemented with different lysine concentrations ($x$-axis). Cells per mL have been deduced from CFU counts. (b-c) Growth curves via OD measurements taken every ∼15 minutes. Plots for the (b) susceptible and (c) resistant strains indicate that lysine has a negligible impact on the exponential growth rates of both strains. Resistant cells supplemented with low amounts of lysine reach lower capacities. Without lysine resistant cells do not grow, while susceptible cells grow to similar carrying capacities with or without lysine.

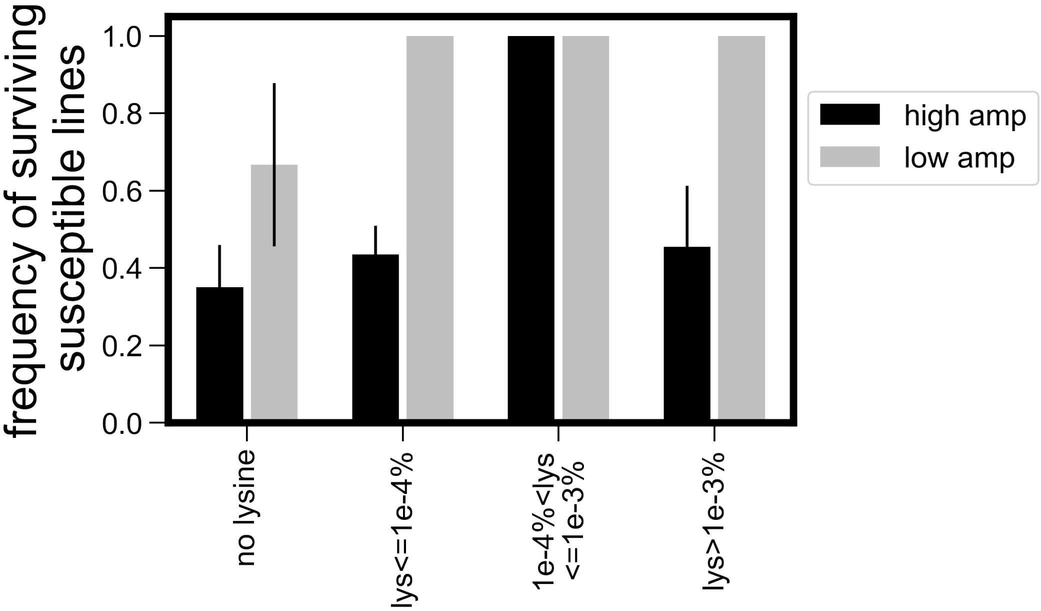


**Figure S2. Survival frequencies of susceptible population in different lysine and antibiotic concentrations.** The average frequency at which the susceptible population survives when evolved in co-culture with the resistant strain, as a function of lysine concentration (*x*-axis) and at high ($\geq 64\mu$g/mL, black) as well as low ampicillin concentrations ($< 64\mu$g/mL, grey). Error bars represent s.e.m.

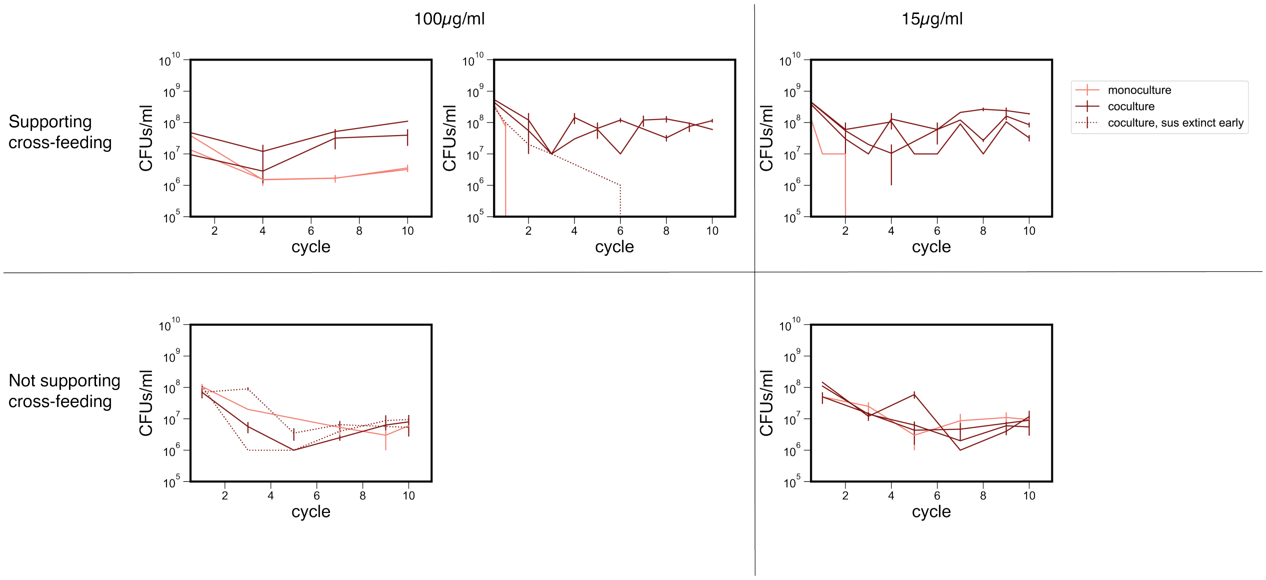


**Figure S3. Evidence for susceptible strain being weakly beneficial to the resistant strain.** Comparison of the resistant strain cell counts at the end of each growth cycle from either mono-cultures (pink) or co-cultures (dark red), grown with the susceptible strain. Shown are results from several evolutionary experiments performed in the absence of lysine with high (100$\mu$g/mL) and low (15$\mu$g/mL) ampicillin. Three sets show resistant strain reaches higher counts in co-cultures compared to mono-cultures (upper panel). Two other sets show similar counts for the resistant strain in mono and co-cultures. However, in three of the co-cultures the susceptible cells went extinct early (before the sixth cycle, indicated by dashed lines), making them similar to mono-cultures. Thus, not considering lines in which the susceptible cells went extinct, 7 lines show higher counts in co-cultures compared to mono-cultures and only 4 lines show similar counts, with an overall increase in resistant population in co-cultures($6 \times 10^7$ cells/mL) compared to mono-cultures ($4 \times 10^6$ cells/mL); $P < 0.005$, two-sided Mann-Whitney U-test. These results support the assumption the susceptible strain is weakly beneficial to the resistant strain.

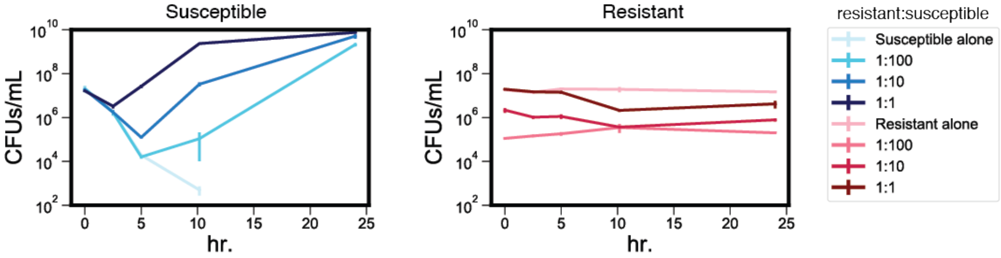


**Figure S4. Initial resistant cell density affects susceptible strain dynamics and final abundances over a growth cycle.** Cell densities of the susceptible (left) and resistant strains over a single growth cycle (time in hr). Overnight samples of susceptible and resistant strains were inoculated either alone or at different ratios of resistant and susceptible cells, into media containing 100$\mu$g/mL ampicillin and no supplemented lysine. For each strain we monitored its abundance by plating on selective plates at different time points during a 24 hr cycle. Each cell density represents the mean of at least 2 replicates. Error bars represent s.e.m.

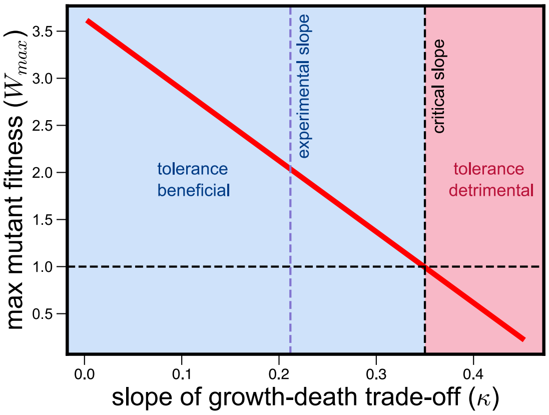


**Figure S5. For tolerance to be beneficial, the growth-death trade-off must be less steep than a critical value.** The maximum fitness of a tolerant mutant $W_{max}$ as a function of the slope $\kappa$ of the trade-off between death rate and growth rate of all evolved tolerant strains ($\kappa_{obs} \approx 0.21$). Beyond a critical slope ($\kappa \approx 0.35$), all tolerant strains are less fit than their susceptible ancestor, and tolerance is no longer beneficial. These calculations were done for no lysine conditions, i.e., when $R_{sat} = 1.2 \times 10^7$ cells.

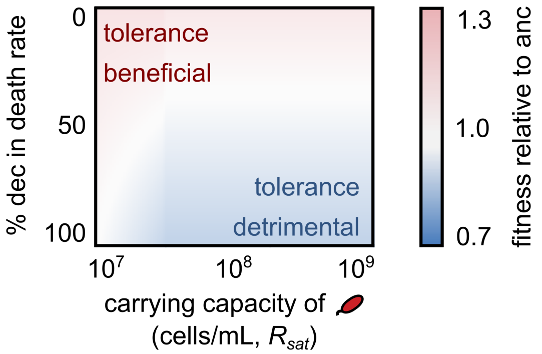


**Figure S6. Our model predicts that tolerant strains will have a lower decrease in death rate (degree of tolerance) in low antibiotic concentrations.** Phase diagram of possible tolerant mutants, similar to Fig. 2f, except for a lower antibiotic concentration (15 $\mu$g/mL). Here, $\kappa = 0.21$, similar to what is experimentally observed. Red represents conditions where tolerant strains are fitter (relative fitness greater than 1), blue represents conditions where tolerant strains are less fit, and white represents conditions where tolerance is neither beneficial nor detrimental. In these conditions, a much lower reduction in death rate (compared with high antibiotic concentrations of 100 $\mu$g/mL shown in Fig. 2f) is predicted to evolve. This is consistent with our experimental observations, shown in Fig. S10.

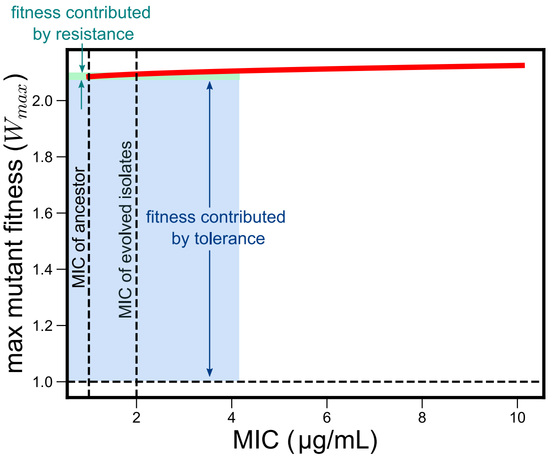


**Figure S7. The maximum fitness of a tolerant strain depends weakly on its MIC in the model.** The maximum relative fitness $W_{max}$ of a tolerant mutant, in conditions of low $R_{sat}$ in the model, as a function of its MIC. The vertical dashed line represents the MIC of the susceptible strain in the model, and the horizontal dashed line represents $W = 1$, the relative fitness of the susceptible strain. Increasing the MIC increases the fitness of a tolerant strain, but the dominant contribution comes from its reduced death and growth rates (its tolerance), not its MIC.

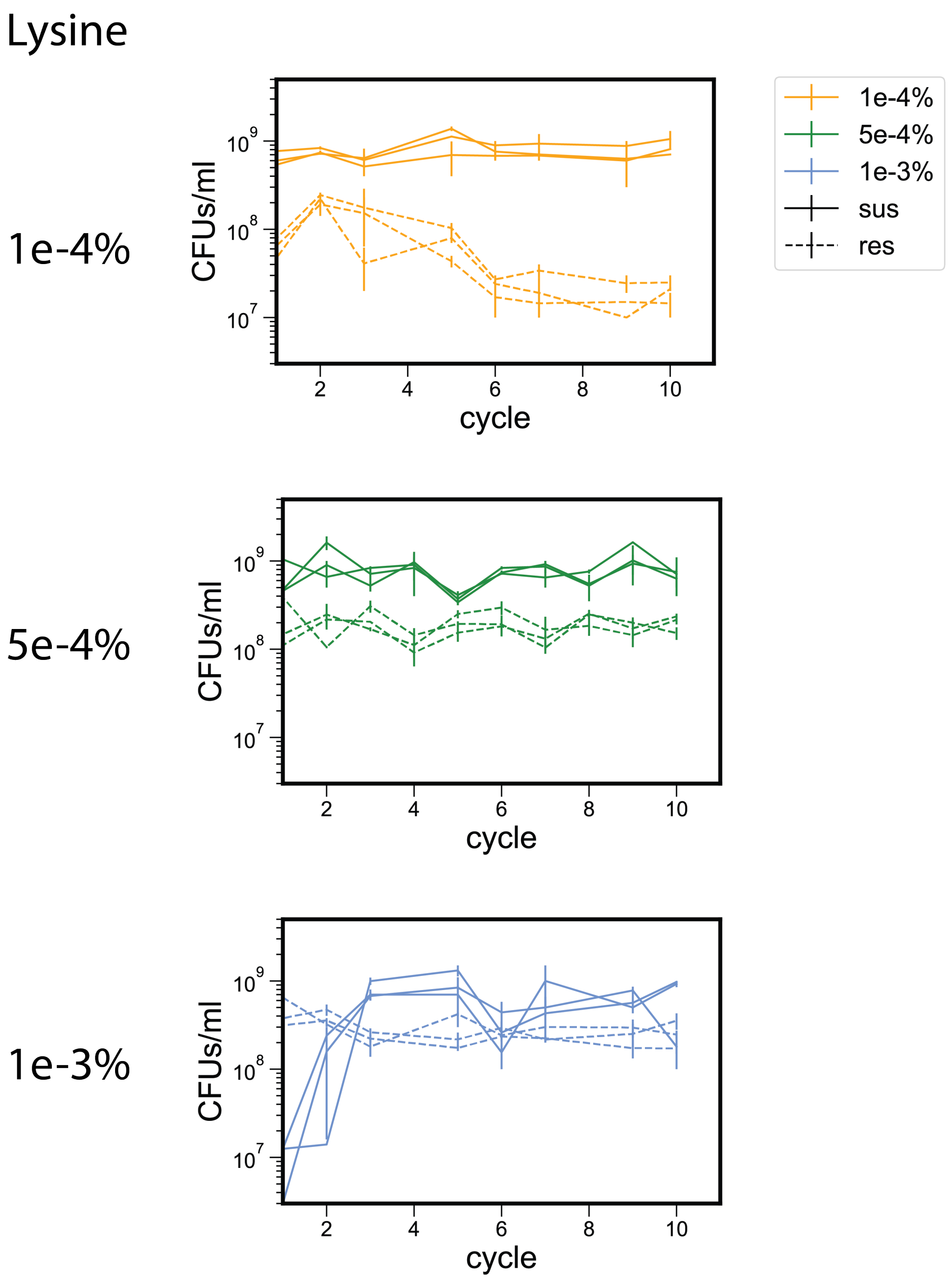


**Figure S8. Resistant carrying capacity increases with increasing lysine concentration.** Cell densities of the resistant auxotroph strain (dotted lines) and susceptible (solid lines) over successive growth cycles, in conditions with different lysine concentrations supplemented. For the data shown, communities containing both susceptible and resistant strains were evolved in media containing 100 $\mu$g/mL ampicillin and the lysine concentrations shown (left of each panel; $n = 3$ for each concentration). Strain abundances were monitored using plating and CFU counting at the end of each cycle. Each point represents the mean of multiple technical repeats, and the error bars represent s.e.m.

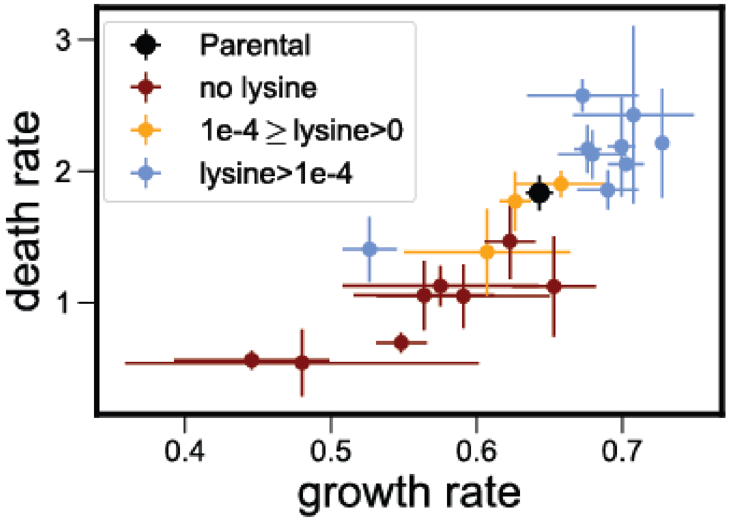


**Figure S9. Increasing lysine concentration lowers the degree of tolerance evolved in susceptible populations.** Death and growth rates of the susceptible ancestor ("parental") and evolved isolates at high ampicillin concentrations (64–100 $\mu$g/mL), colored by the lysine concentration they were evolved in. Each point represents the mean , whereas error bars represent the s.e.m.

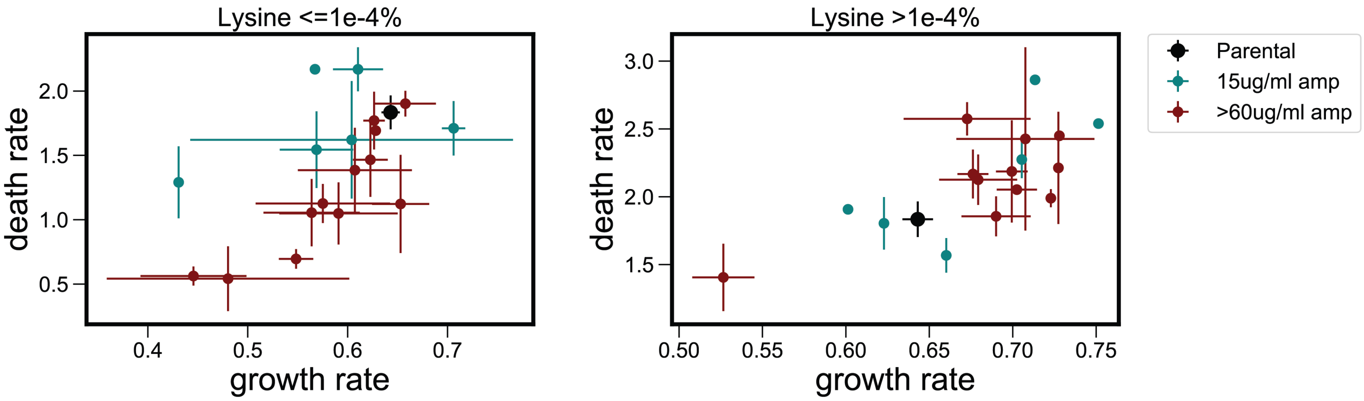


**Figure S10. Lowering ampicillin concentration lowers the degree of tolerance evolved in susceptible populations.** Death and growth rates of the susceptible ancestor ("parental" colored black) and evolved isolates at different ampicillin concentrations (15$\mu$g/mL blue, $64 - 100\mu$g/mL dark red ) and in different lysine concentrations (lower concentrations, less than $10^{-4}$%, on the left, higher concentrations on the right). Under low lysine, lowering ampicillin concentrations showed a significant difference in death rate; Mann Whitney one-sided test $P < 0.05$ while no significant change in death rates was measured between high and low ampicillin under high lysine conditions; Mann Whitney one-sided test $P > 0.1$ Each point represents the mean , whereas error bars represent s.e.m. The decreased tolerance as a function of ampicillin concentration is only observed for lower concentrations of lysine.

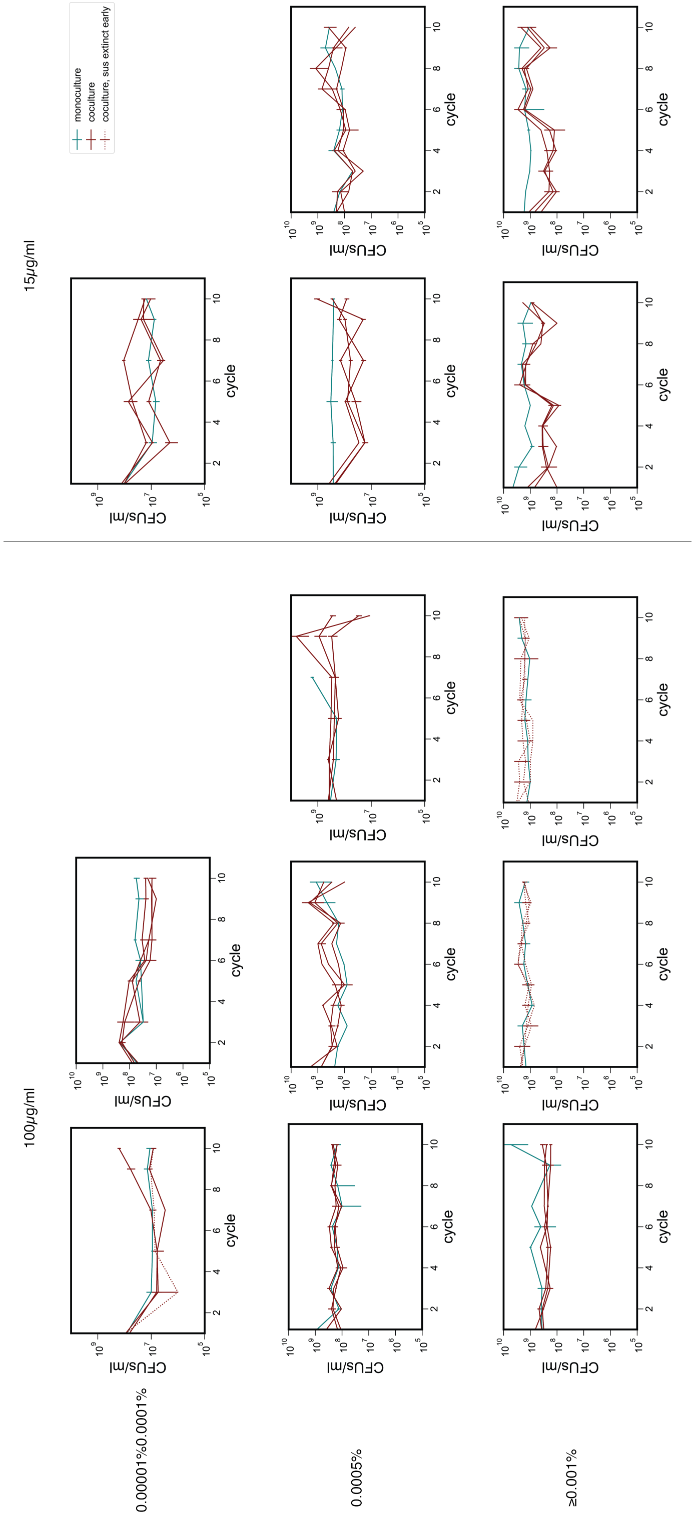


**Figure S11. Susceptible strain is not beneficial to resistant strain when lysine is added** Comparison of the resistant strain counts at the end of cycles from mono-cultures (blue) and co-cultures (dark red) grown with the susceptible strain. Different evolutionary experiments carried with different concentrations of lysine carried in high (100$\mu$g/mL) and low (15$\mu$g/mL) ampicillin. Co-cultures do not reach higher counts suggesting when lysine is present the resistant strain does not benefit from the presence of the susceptible strain. The capacity of the resistant cells do tend to increase when more lysine is supplied.

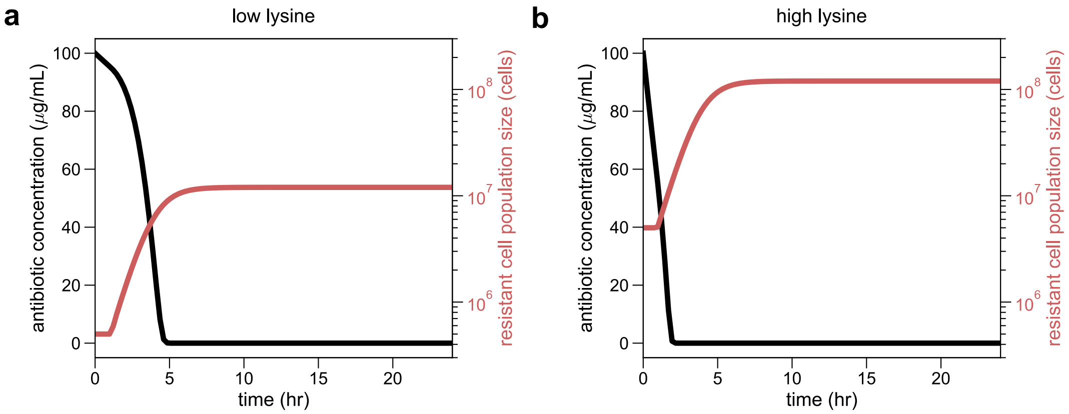


**Figure S12. Dynamics of antibiotic degradation simulated using our mathematical model.** Time series of antibiotic concentration (black) and resistant cell population size (red) as simulated according to our mathematical model (for details, see Methods; for parameters, see table S1 of the Supplementary Materials), in (a) low (or no) lysine conditions, where $R_{sat} = 1.2 \times 10^7$ cells, and (b) high lysine conditions, where $R_{sat} = 1.2 \times 10^8$ cells, cartoons of which were shown in Fig. 2b and Fig. 2d, respectively.
